# Application of organic fertilisation such as poultry manure enhanced carbohydrate, fat and vitamin C contents of Huckleberry (*Solanum scabrum*) more than mineral fertilisation in Buea-Cameroon

**DOI:** 10.64898/2026.10.30.763013

**Authors:** Kwanda Belinda Nuigho, Schneideur Ndongho Tanting, Germain Sotoing Taiwe, Tange Denis Achiri, Gillian Nkeudem Asoba

## Abstract

Huckleberry (*Solanum scabrum*), is a vital indigenous leafy vegetable in sub-Saharan Africa(SSA) and Cameroon, serving as a rich source of vitamins, proteins, and minerals. This study evaluated the nutritional composition, growth, and yield parameters of huckleberry produced using organic and inorganic fertilizers in Buea. A randomized complete block experiment with four blocks compared four fertilisation regimes: pig manure, poultry manure, control, and urea. 6 weeks after planting (WAP), growth parameters; plant height, leaf number and leaf area were measured alongside cumulative yield. Harvested samples were analysed to determine total protein, carbohydrate, fat, and vitamin C contents. Data were subjected to analyse using a two-way analysis of variance. Fertilizer application significantly (*P*<0.05) enhanced plant performance compared to the unfertilized control. At 8 WAP, urea had the most vigorous vegetative growth, having the highest mean plant height (25.92 cm), maximum leaf count (29.38), and total yield (20.72 tons/ha). Nutritionally, urea produced the highest total protein content (290.30 mg/100 g), but reduced vitamin C level (24.20 mg/100 g) below the baseline of the control (27.53 mg/100 g). Poultry manure had the highest content of carbohydrates (6.52%), fats (7.8%), and vitamin C (42.57 mg/100 g). The study demonstrates a clear trade-off between crop yield and specific nutritional attributes based on nutrient sources. While urea enhanced plant growth and yield, organic amendments like poultry manure improved the micronutrients like carbohydrate and fat composition of huckleberries. Future work should focus on integrated application of poultry and pig manure with urea to see their effects of huckleberry performance, nutrient contents and soil health.

## 1. Introduction

Huckleberry (*Solanum scabrum*), also called African nightshade and locally known as country *njama njama* is a green, indigenous leafy vegetable that grows around the world [1], its utilization can deliver more nutrients and phytochemicals into the diet. In Cameroon, it is used in the preparation of traditional delicacies or served as a side dish such as in Kati-kati, and achu. It is a rich source of vitamins, minerals, protein, fat, carbohydrate, fibre, β-carotene, and ascorbic [2]. Plant cultivation and processing may impact nutrient and phytochemical content of vegetables [3]. The cultivation of huckleberry, like many other crops, is significantly influenced by nutrient availability in the soil, which can be manipulated through the application of fertilizers. The nutrient values of traditional leafy vegetables may vary with soil fertility, plant age and the type [4]. Fertilizers are broadly categorized into organic and inorganic [5], and studies done by [4, 6–7], have shown fertilizers to significantly influence crop growth and yield and ultimately, the nutritional composition of the harvested produce. Organic fertilizer (manure), derived from natural sources such as compost, manure, kitchen waste and plant residues, are known for improving soil structure, enhancing microbial activity, and providing a slow release of nutrients [7]. In contrast, inorganic fertilizers, synthetically produced [8], offer readily available nutrients in precise concentrations, often leading to rapid plant growth. While the general benefits of huckleberries are well-documented [9], there is a rareness of inclusive, comparative study specifically examining the influence of organic versus inorganic fertilizers on their detailed nutritive composition. The majority of existing literature focuses on yield, general plant health or growth, with less importance on the changes in micronutrient, vitamin, and phytochemical composition Understanding the differential effects of these fertilizer types on the nutritional profile of huckleberries is important for enhancing cultivation practices to boost health-promoting qualities vegetables. This work aims to compare the nutritional content of huckleberries grown under the influence of organic and inorganic fertilizers. We hypothesised that: (i) organic and inorganic fertilisers applications will enhance huckleberry growth and yield; (ii) organic fertiliser like poultry manure will improve huckleberry fat, vitamin C and carbohydrate contents

## 2. Materials and methods

### 2.1. Experimental site and design

This study was conducted from February to May 2026 at Lysoka village, Buea, Southwest Region of Cameroon (Latitude: 4°08′55″ N, Longitude: 9°17′01″ E). The site lies within Cameroon’s humid forest agroecological zone (Zone IV). Field experiment was established in a randomized complete block design. The field was divided into four blocks with each block representing a replicate and each experimental unit measured 3m x3m, giving 16 experimental units. The inter-block spacing was 1.5 m while the distance in between two plots was 1 m with a border of 1m round the plot used. The entire 400 m^2^ experimental field was cleared, raked and laid out using a measuring tape, ropes and a stick to mark the various plots for easy identification during ploughing.

### 2.2. Field preparation

#### 2.2.1. Nursery preparation

Huckleberry seeds (Foumbot variety) were gotten from Agricultural Research Institute for Development (IRAD), Foumbot. Nursery beds were prepared, grasses removed and beds of 1 x 1 m were raised and the seed sown by broadcasting on the beds and covered with light soil. However, before sowing the beds were treated with insecticide and fungicide to prevent insects from eating up the seeds and also, to limit the growth of fungi. Dry hay was used as the mulching material for 3 -5 days and was later removed. Watering was done with the use of a watering can that has the shower head attached to it. This process was carried out early in the morning and at sunset 5- 6pm in the evening and was transplanted to the field 3 weeks after planting.

#### 2.2.1. Transplanting, treatments application and field maintenance

A total of 4 treatments were used to investigate the effect of fertilisers application on huckleberry performance and nutrients content in the various plots. These treatments were pig manure, poultry manure, control, and urea (Table 1). Each of the above treatments were randomized on 4 different plots within the various blocks. For the control plot, they were no application of fertilisation. Urea (purchased from a registered agro-dealer in Buea was, and applied at 100 kg product ha^-1^ (equivalent to 46 kg N ha^-1^) in two equal splits at at 3 days after transplanting (DAT) and the second at 14 DAT. Poultry manure was obtained from broiler farm at the, Faculty of Agriculture and Veterinary Medicine, University of Buea, and applied at 8 tons ha^-1^ oven-dry weight basis (12.8 kg per 16 m^2^ plot), broadcast and incorporated two weeks before transplanting ; the manure contained 100 % dry matter, 2.5 % total N, 2.0 % P, 0.87 % K, 28 % organic carbon and a C:N ratio of 8, corresponding to 280 kg N ha^-1^.Pig manure, obtained from small holder farmers farm in Buea and applied at 8 tons ha^-1^ oven-dry weight basis (12.8 kg per 16 m^2^ plot), broadcast and incorporated two weeks before transplanting ; the manure contained 100 % dry matter, 2.3 % total N, 1.8 % P, 0.80 % K, 25 % organic carbon and a C:N ratio of 7, corresponding to 270 kg N ha-1.Transplanting was done at seedling age of three weeks after nursing and was carried out in the evening. The seedlings were planted right to the first leaves using the planting distance of 20 x 20 cm giving a plant density of 25 plants / m^2^ with a total of 225 plants per experimental unit, and watering exercise was done after the transplanting process.

**Table 1:** Treatments·of the experiment and. their d1eserip1tion

| **Treatments** | **Description** |
|---|---|
| T1 | Pig manure |
| T2 | Poultry manure |
| T3 | Control |
| T4 | Urea |

To ensure optimal growth and yield of huckleberry (*Solanum scabrum*), manual weeding was conducted as needed using hoe to remove competing weeds. Weeding was performed carefully around huckleberry plants to avoid root disturbance, with major session typically occurring at week two after transplanting at the critical weed period of huckleberry [10], in order to prevent the weeds from competing with the crops. The crop was rain-fed throughout, other pests like snails were hand-picked and crushed.

### 2.3. Data collection

#### 2.3.1. Growth parameters and fresh weight

Growth data were recorded at 1,2 and 3 weeks after transplanting (WAT) on the same 10 tagged plants in the two central rows of each plot; the mean of the 10 plants was used as the plot value, so that the plot (n = 4 per treatment) was the unit of analysis. The total number of leaves were counted on each plant; this was done on the 10 tagged plants. Vegetable height was measured with a measuring tape from the base of the plant to the tip of the youngest leaf (cm). The leaf length of the vegetable plant was measured with a tape from the base of the leaf to the tip of the leaf and the leaf width (cm) was also measured, individual leaf area (cm^2^) was calculated as leaf length x maximum leaf width.

At maturity 4 WAT, harvesting was carried out with the use of a sharp knife early in the morning. Harvesting was carried out within a one square meter quadrat per treatment plot and was done for 4,5,6 and 7 WAT and total yield calculated per treatment. Harvested plants were cut with a sharp knife and Fresh weight was obtained using a digital electric balance recorded in Kg and converted to tonsha^-1^.

#### 2.3.2. Nutritional analysis

Fresh huckleberry harvested at 3WAT was taken for nutritional analysis at the Agro ecology laboratory at the University of Buea, Faculty of Agriculture and Veterinary Medicine laboratory and evaluated for total protein, total carbohydrates, vitamin C, and fat content. Fat content was determined using the Soxhlet Fat Extraction Method AOAC. Protein content examined using the AOAC’s Kjehdal technique. Carbohydrate content was analyzed using the enzyme method, and Vitamin C content determined using 2,6-Dichlorophenol Indophenol AOAC method [11].

### 2.4. Data analysis

All analyses were carried out in IBM SPSS Statistics version 23 on plot means (n = 4 plots per treatment). Growth data recorded at 1,2 and 3 WAT were analysed. Normality of model residuals was checked with the Shapiro-Wilk test and homogeneity of variances with Levene’s test; no transformation was required before analysis. To test the effects of the different fertiliser regimes, treatment effects were tested by analysis of variance and, means were separated with Tukey’s HSD test at P < 0.05. This analysis was also conducted for vegetable growth parameters (Vegetable height, leaf area and number of leaves), fresh weight and nutrients content.

## 3. Results

### 3.1. Growth parameters

At 1 WAT, vegetable height differed significantly among treatments (*P*<0.05, Table 2). The value ranges from 9.17-13.52 cm, with the highest in the urea (13.52 cm) treatment, followed by poultry manure (12.17 cm) and pig manure (9.91 cm) and lowest in the control(9.17cm). Application of urea, poultry manure and pig manure significantly increased vegetable height by 47.44 %,32.72 % and 8.07 % respectively. At 2 WAT, vegetable height differed significantly among treatments (*P*<0.05, Table 2). The value ranges from 15.36-19.72 cm, with the highest in the urea (19.72 cm) treatment, followed by poultry manure (18.37 cm) and pig manure (16.09 cm) and lowest in the control(15.36cm). Application of urea, poultry manure and pig manure significantly increased vegetable height by 28.39 %,19.60 % and 4.75 % respectively. At 3 WAT, vegetable height differed significantly among treatments (*P*<0.05, Table 2). The value ranges from 19.89-25.92 cm, with the highest in the Urea (25.92 cm) treatment, followed by poultry manure (24.66 cm) and pig manure (21.68 cm) and lowest in the control(19.89cm). Application of urea, poultry manure and pig manure significantly increased vegetable height by 30.32 %,23.89 % and 9.00 % respectively.

**Table 2:** Effects of fertilisation on huckleberry height (cm)

| **Treatment** | **1 WAT** | **2WAT** | **3WAT** |
|---|---|---|---|
| Pig manure | 9.91 ± 1.70a | 16.09 ± 2.00a | 21.68 ± 2.73b |
| Poultry manure | 12.17 ± 2.28b | 18.37 ± 2.28b | 24.66 ± 2.58c |
| Control | 9.17 ± 1.10a | 15.36 ± 1.08a | 19.89 ± 3.52a |
| Urea | 13.52 ± 2.03c | 19.72 ± 2.03c | 25.92 ± 2.03c |

Bars (Mean ± SD) within column with different letters between treatments in the same week are significantly different (Tukey’s HSD, *P*<0.05).

At 1 WAT, number of leaves differed significantly among treatments (*P*<0.05, Table 3). The value ranged from 8.58-16.75, with the highest in the urea (16.75) treatment, followed by poultry manure (13.29) and control (8.83) and lowest in pig manure (8.58). Application of urea and poultry manure significantly increased vegetable number of leaves by 89.69 %, and 50.51 % respectively when compared to the control treatment. At 2 WAT, number of leaves differed significantly among treatments (*P*<0.05, Table 3). The value ranged from 14.70-22.79, with the highest in the urea (22.79) treatment, followed by poultry manure (19.29) and control (14.83) and lowest in pig manure (14.70). Application of urea and poultry manure significantly increased vegetable number of leaves by 53.67 %, and 30.07 % respectively when compared to the control treatment. At 3 WAT, number of leaves differed significantly among treatments (*P*<0.05, Table 3). The value ranged from 21.66-29.38, with the highest in the Urea (29.38) treatment, followed by poultry manure (26.29) and control (21.92) and lowest in pig manure (21.66). Application of urea and poultry manure significantly increased vegetable number of leaves by 34.03 %, and 19.94 % respectively when compared to the control treatment.

**Table 3:**
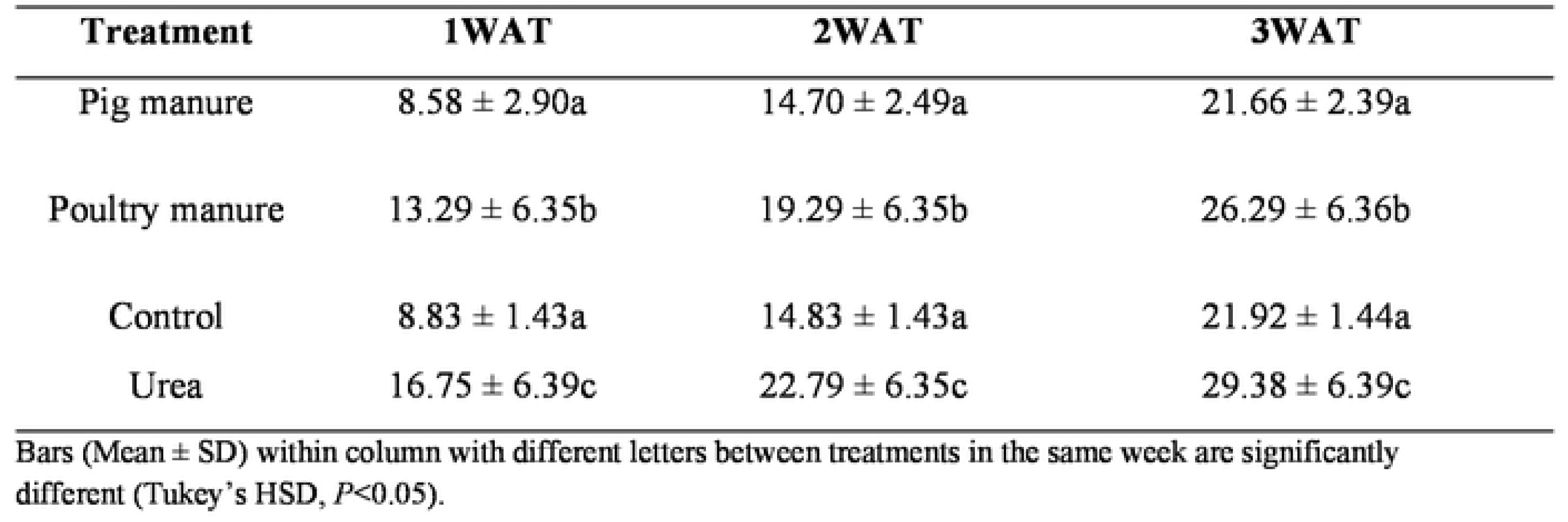
Effects of fertilisation on huckleberry number of leaves

| **Treatment** | **1WAT** | **2WAT** | **3WAT** |
|---|---|---|---|
| Pig manure | 8.58 ± 2.90a | 14.70 ± 2.49a | 21.66 ± 2.39a |
| Poultry manure | 13.29 ± 6.35b | 19.29 ± 6.35b | 26.29 ± 6.36b |
| Control | 8.83 ± 1.43a | 14.83 ± 1.43a | 21.92 ± 1.44a |
| Urea | 16.75 ± 6.39c | 22.79 ± 6.35c | 29.38 ± 6.39c |

Bars (Mean ± SD) within column with different letters between treatments in the same week are significantly different (Tukey’s HSD, *P*<0.05).

At 1 WAT, leaf area differed significantly among treatments (*P*<0.05, Table 4). The value ranged from 29.22-32.67 cm^2^, with the highest in the urea (32.67 cm^2^) treatment, followed by poultry manure (31.49 cm^2^) and pig manure (29.44 cm^2^) and lowest in control (29.22 cm^2^). Application of urea, poultry manure and pig manures increased vegetable leaf area by 11.81 %, 7.77 % and 0.75 % respectively when compared to the control treatment. At 2 WAT, leaf area differed significantly among treatments (*P*<0.05, Table 4). The value ranged from 55.82-60.89 cm^2^, with the highest in the urea (60.89 cm^2^) treatment, followed by poultry manure (59.36 cm^2^) and control (55.82 cm^2^) and lowest in pig manure (52.99 cm^2^). Application of urea and poultry manure increased vegetable leaf area by 9.08 % and 6.34 % respectively when compared to the control treatment. At 3 WAT, leaf area differed significantly among treatments (*P*<0.05, Table 4). The value ranged from 81.49-97.61 cm^2^, with the highest in the urea (97.61 cm^2^) treatment, followed by poultry manure (96.06 cm^2^) and control (90.02 cm^2^) and lowest in pig manure (81.49 cm^2^). Application of urea and poultry manure increased vegetable leaf area by 8.43 % and 6.71 % respectively when compared to the control treatment.

**Table 4:**
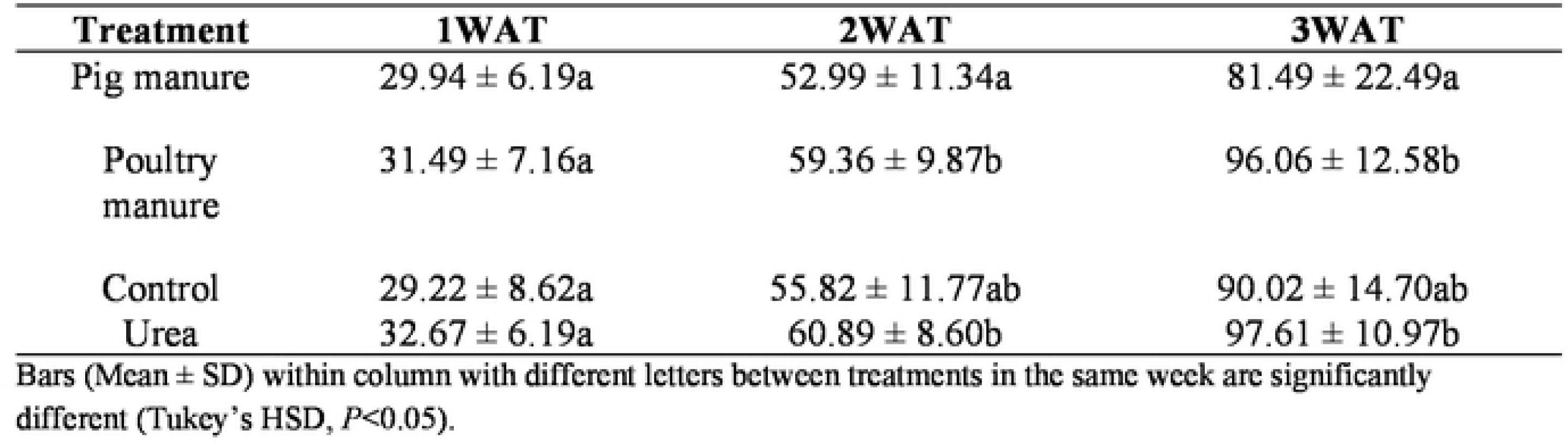
Effect of fertilisa1Jon on huckleberry leaf area (cm^2^)

| **Treatment** | **1WAT** | **2WAT** | **3WAT** |
|---|---|---|---|
| Pig manure | 29.94 ± 6.19a | 52.99 ± 11.34a | 81.49 ± 22.49a |
| Poultry manure | 31.49 ± 7.16a | 59.36 ± 9.87b | 96.06 ± 12.58b |
| Control | 29.22 ± 8.62a | 55.82 ± 11.77ab | 90.02 ± 14.70ab |
| Urea | 32.67 ± 6.19a | 60.89 ± 8.60b | 97.61 ± 10.97b |

Bars (Mean ± SD) within column with different letters between treatments in the same week are significantly different (Tukey’s HSD, *P*<0.05).

### 3.2. Fresh weight

At 4 WAT, fresh weight differed significantly among treatments (*P*<0.05, Table 5). The value ranges from 1.09-3.1 tonsha^-1^, with the highest in the urea (3.1 tonsha^-1^) treatment, followed by poultry manure (1.9 tonsha^-1^) and pig manure (1.25 tonsha^-1^) and lowest in the control (1.09 tonsha^-1^). Application of urea, poultry manure and pig manure increased fresh weight by 184.4 %, 74.31 % and 14.68 % respectively when compared to the control. At 5 WAT, fresh weight differed significantly among treatments (*P*<0.05, Table 5). The value ranges from 0.9-4.5 tonsha^-1^, with the highest in the urea (4.5 tonsha^-1^) treatment, followed by poultry manure (2.9 tonsha^-1^) and joint lowest in pig manure (0.9 tonsha^-1^) and control (0.9 tonsha^-1^). Application of urea and poultry manure increased fresh weight by 400 % and 222.22 % respectively when compared to the control. At 6 WAT, fresh weight differed significantly among treatments (*P*<0.05, Table 5). The value ranges from 1.56-6.09 tonsha^-1^, with the highest in the urea (6.09 tonsha^-1^) treatment, followed by poultry manure (4.5 tonsha^-1^) and pig manure (2.03 tonsha^-1^) and lowest in the control (1.56 tonsha^-1^). Application of urea, poultry manure and pig manure increased fresh weight by 290.38 %, 188.46 % and 30.13 % respectively when compared to the control. At 7 WAT, fresh weight differed significantly among treatments (*P*<0.05, Table 5). The value ranges from 2.5-7.03 tonsha^-1^, with the highest in the urea (7.03 tonsha^-1^) treatment, followed by poultry manure (5.6 tonsha^-1^) and pig manure (2.96 tonsha^-1^) and lowest in the control (2.5 tonsha^-1^). Application of urea, poultry manure and pig manure increased fresh weight by 181.2 %, 124 % and 18.40 % respectively when compared to the control. Total fresh weight differed significantly among treatments (*P*<0.05, Table 5). The value ranges from 6.05-20.72 tonsha^-1^, with the highest in the urea (20.72 tonsha^-1^) treatment, followed by poultry manure (14.9 tonsha^-1^) and pig manure (7.14 tonsha^-1^) and lowest in the control (6.05 tonsha^- 1^). Application of urea, poultry manure and pig manure increased total fresh weight by 242.48%, 146.28 % and 18.02% respectively when compared to the control.

**Table 5:**
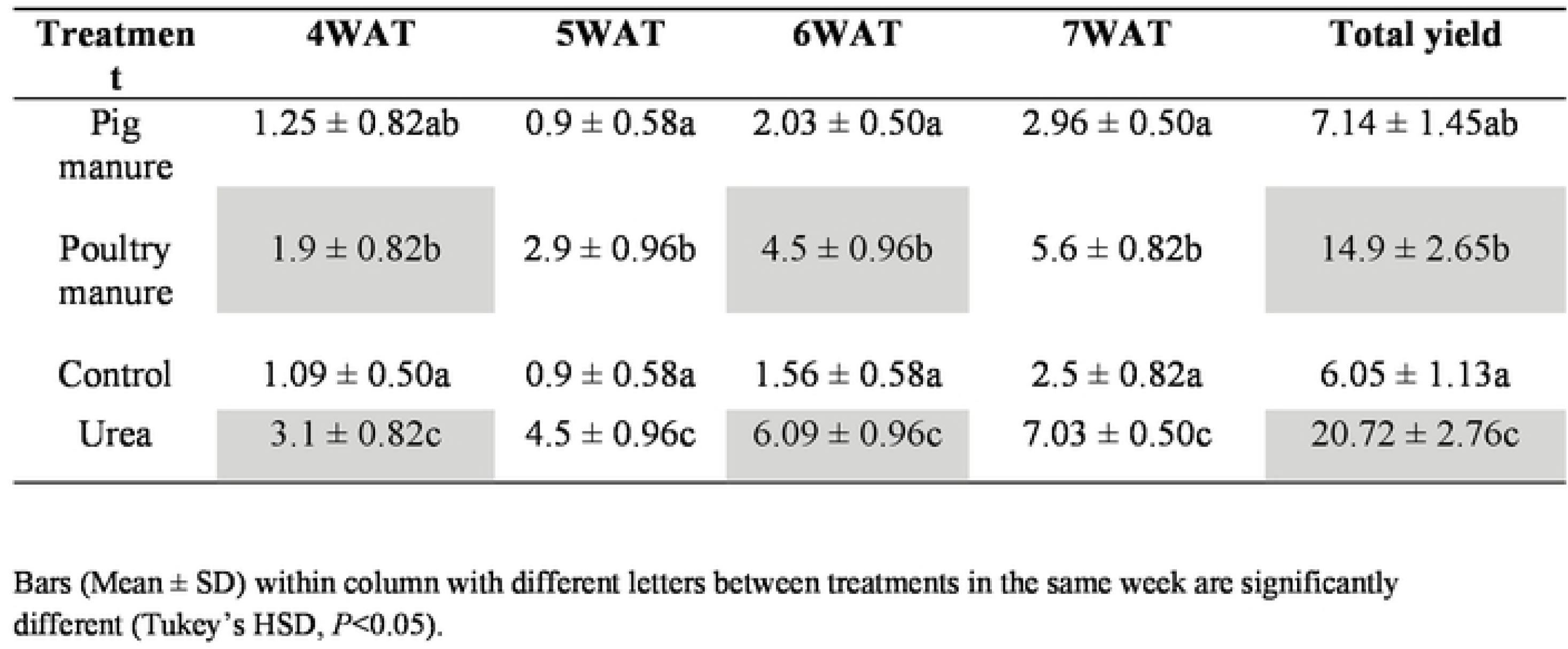
Effects of fertilisation on huckleberry yield(tons/ha)

### 3.3. Nutritional contents

Carbohydrate content differed significantly among treatments (*P*<0.05, Table 6). The value ranged from 4.25-6.52 %, with the highest in poultry manure (6.52 %) treatment, followed by pig manure (5.92 %) and urea (5.25 %) and lowest in the control (4.25 %). Application of poultry manure, pig manure and urea increased carbohydrate content by 53.41 %, 39.29 % and 23.53 % respectively when compared to the control. Protein content differed significantly among treatments (*P*<0.05, Table 6). The value ranged from 185-290.3 mg/100 g, with the highest in urea (290.3 mg/100 g) treatment, followed by poultry manure (260.4 mg/100 g) and pig manure (235.1 mg/100 g) and lowest in the control (185 mg/100 g). Application of urea, poultry manure and pig manure increased protein content by 56.92 %, 40.76 % and 27.08 % respectively when compared to the control. Fat content differed significantly among treatments (*P*<0.05, Table 6). The value ranged from 3.43-7.8 %, with the highest in poultry manure (7.8%) treatment, followed by urea (6.57 %) and pig manure (6.0 %) and lowest in the control (3.43%). Application of poultry manure, urea and pig manure increased fat content by 127.41 %, 91.55 % and 74.93 % respectively when compared to the control. Vitamin C content differed significantly among treatments (*P*<0.05, Table 6). The value ranged from 24.20-42.57 mg/100 g, with the highest in poultry manure (42.57 mg/100 g) treatment, followed by pig manure (36.53 mg/100 g) and control (27.53 mg/100 g) and lowest in urea (24.20 mg/100 g).

**Table 6:** Effects. on fertilisation on nutritional composition of huckleberry

| **Treatment** | **Carbohydrates (%)** | **Protein (mg/100 g)** | **Fat (%)** | **Vitamin C (mg/100 g)** |
|---|---|---|---|---|
| Control | 4.25 ± 0.65a | 185.00 ± 2.23a | 3.43 ± 1.20a | 27.53 ± 3.49b |
| Pig manure | 5.92 ± 0.61c | 235.10 ± 30.03b | 6.00 ± 1.54b | 36.53 ± 2.53c |
| Poultry manure | 6.52 ± 0.71c | 260.40 ± 40.37c | 7.8 ± 0.95d | 42.57 ± 2.56d |
| Urea | 5.25 ± 0.81b | 290.30 ± 13.85d | 6.57 ± 1.55c | 24.20 ± 2.22a |

Bars (Mean ± SD) within column with different letters between treatments in the same week are significantly different (Tukey’s HSD, *P*<0.05).

Application of poultry manure and pig manure increased vitamin C content by 54.57 %, and 32.69 % respectively when compared to the control.

## 4. Discussion

### 4.1. Effect of fertilisation on growth and yield of huckleberry

This study found that the fertiliser treatments had a significant effect on huckleberry growth and yield. Overall, the data show that fertiliser application outperforms the control, indicating the importance of nitrogen supplementation for crop development. Among the treatments examined, urea consistently provided greater vegetative growth and yield, but poultry manure exhibited relatively higher effects on some quality parameters. Specifically, when it came to plant height and number of leaves, urea consistently provided the most noticeable vegetative response. This result is in line with [12], demonstrating that nitrogen fertilisation improves *Solanum scabrum* and other leafy vegetable growth and yield. The significant performance of urea treatment in this study is most likely due to the rapid availability of nitrogen in mineral form, which may have encouraged early and sustained vegetative growth more effectively than organic sources. This is consistent with the findings of [13], who observed that combined applications of farmyard manure and calcium ammonium nitrate improved plant height and yield parameters of African nightshade, demonstrating that nitrogen supply is a primary factor of growth in Solanum species.

Furthermore, poultry manure also generated significant improvements, especially in early leaf formation and leaf area development, even though urea was the most effective treatment for overall vegetative growth. Poultry manure outperformed pig manure and the control in the first week of leaf emergence, suggesting that organic additions may promote early establishment through improved soil physical conditions and slow nutrient release. The manure from poultry significantly increased vegetative growth and yield. [14], reported similar findings, noting that *Solanum macrocarpon* growth and leaf area were enhanced by both organic and inorganic fertilisers, however the degree of responses differed depending on the kind of fertiliser. Similarly, [15] discovered that *Solanum scabrum* growth characteristics were much improved by fortified liquid organic manure, suggesting that organic nutrient sources might encourage rapid growth when nutrient release is timed to match crop need. According to the current findings, poultry manure may promote more slow and balanced vegetative development whereas urea offers a higher rapid growth response. More so, yield response followed the same pattern as vegetative growth, with urea being the most productive, followed by poultry manure, and pig manure. This finding identified yield as the most relevant cost- effective indicator, and urea’s performance in this regard emphasises the necessity of easily readily available nitrogen for marketable leafy vegetable crop. The result of this finding is consistent with previous observations that mineral nitrogen sources can significantly boost production in leafy vegetables, especially where soil nutrient availability is limited [12–13]. In those studies, leafy vegetable yield increased significantly with mineral nitrogen or integrated fertiliser application, supporting the notion that leafy indigenous fruits and veggies respond well to nutrients and mineral additions. The significantly lower production under pig dung in this study could be related to changes in nutrient composition, mineralisation rate, or nutrient alignment with crop uptake. This highlights the fact that not all organic manures, even when applied at comparable amounts, could have the same economic effects. This results therefore support our hypothesis that application of organic and inorganic amendments will increase huckleberry growth and yield.

### 4.2. The effects of organic manure and inorganic fertiliser on nutritional composition of huckleberry

Urea had the highest protein concentration, reaching 290.30 mg/100 g, showing that nitrogen plays an important role in amino acid and protein synthesis. This finding was in line with [14]. indicating that inorganic nitrogen fertilisation increases protein-related composition in leafy vegetables. The same treatment, however, decreased vitamin C concentration below the control level, suggesting that high nitrogen supply promote protein formation at the expense of other antioxidant molecules. [16–17] showed that whereas mineral fertiliser may increase nitrate accumulation and lower ascorbic acid levels, organic fertility regimes frequently improve vitamin C in leafy vegetables. Poultry manure had the highest vitamin C concentration in this study, suggesting that organic nourishment sources might better enhance the synthesis or preservation of specific micronutrients which could be attributed to a more balanced nutrient availability, and slower nutrient release [18]. Additionally, this study’s finding that poultry manure enhanced the contents of fat, vitamin C and carbohydrates more than urea which it implies that organic manure might enhance huckleberry nutritional quality even when production is not maximised [16–17,19]. This therefore support the hypothesis that application of poultry manure will increase huckleberry fat, vitamin C and carbohydrate contents.

## 5. Conclusions

The findings confirm that managing fertilizer for huckleberry should look beyond simple yield maximization. Choosing the right fertilizer requires balancing soil sustainability, production goals, and the nutritional needs of the consumer. The qualitative benefits of poultry manure show the relevance of organic amendments in enhancing food composition and possibly maintaining soil fertility. The dominance of urea for growth and yield underscores the significance of instantly accessible nitrogen in leafy vegetable production. Therefore, the most balanced method would be an amalgamation nutrient management strategy that preserves or enhances the nutritional quality associated with organic inputs while utilising the quick growth response associated with mineral fertiliser. Future research should quantify the long term effects of these practices on vegetable performance and nutritional contents across multiple seasons and climatic conditions before recommendations are made.

## Author contributions

All authors contributed to the study concept. Material preparations, data collections and analyses were performed by Kwanda Belinda Nuigho , Schneideur Ndongho Tanting , Germain Sotoing Taiwe , Tange Denis Achiri and Gillian Nkeudem Asoba. Supervision was done by Gillian Nkeudem Asoba. The first draft of the manuscript was written by Kwanda Belinda Nuigho and Schneideur Ndongho Tanting and all authors commented on previous versions of the manuscript. All authors read and approved the final manuscript.

## Acknowledgements

We acknowledge the Agro ecology Laboratory, Faculty of Agriculture and Veterinary medicine, University of Buea for nutritional analysis.

## Data availability statement

All data generated or analysed during this study are included in this published article

## Declarations Ethics approval

The Huckleberry plant used in this study were cultivated under standard national and international guidelines on agricultural practices and sampled within acceptable agronomic limits on cultivated crops.

## Consent to participate

Huckleberry is economically important, environmentally friendly not threatened by extinction. Therefore, no specific permission or licenses were required for collecting maize plant part.

## Consent to publish

Not applicable

